# Discovery of Resistance Pathways to Fibroblast Growth Factor Receptor inhibition in Bladder Cancer

**DOI:** 10.1101/183293

**Authors:** Sumanta K Pal, Miaoling He, Jeremy O Jones

**Affiliations:** Department of Medical Oncology, City of Hope National Medical Center, Duarte, CA

**Keywords:** NVP-BGJ398, FGFR, bladder cancer, resistance, PIM kinase

## Abstract

**Background:** Aberrant fibroblast growth factor receptor (FGFR) signaling drives the growth of many bladder cancers. NVP-BGJ398 is a small molecule with potent inhibitory activity of FGFRs 1, 2, and 3, and has been shown to selectively inhibit the growth of bladder cancer cell lines that over-express FGFR3 or have oncogenic FGFR3 fusions. As with many agents targeting receptor tyrosine kinases, resistance is known to develop.

**Objective:** We sought to identify potential mechanisms of resistance to NVP-BGJ398 in cell culture models of bladder cancer.

**Methods:** RT-112 bladder cancer cell lines were derived that were resistant to growth in 3uM NVP-BGJ398. RNA-sequencing was performed on resistant and parental cell lines to identify potential resistance mechanisms and molecular experiments were carried out to test these predictions.

**Results:** RNA-seq demonstrated decreased expression of FGFR3 and increased expression of FGFRs 1 and 2 in resistant cell lines. Over-expression of FGFR3 in NVP-BGJ398 resistant cells decreased their proliferation. Pathway analysis of RNA-seq data also implicated PIM kinase signaling, among other pathways, as a potential mediator of resistance. Treatment of BGJ398 resistant cells with the PIM kinase inhibitor SGI-1776 reduced the growth of the cells.

**Conclusions:** Our results suggest that altered FGFR expression and PIM kinase activity could mediate resistance to NVP-BGJ398. These pathways should be investigated in samples from patients resistant to this drug.

## 1. Introduction

Bladder cancer, the vast majority of which is urothelial carcinoma, is the fifth most common cancer and one of the most expensive cancers to treat in the United States due to the length of required treatment and degree of recurrence [1]. Bladder cancers are most readily divided into two major groups depending on the clinical and molecular features; non-muscle invasive and muscle invasive cancers. 70% of cases are diagnosed as non-muscle-invasive bladder cancer (NMIBC) with a favorable prognosis following transurethral resection and intravesical chemotherapy or immunotherapy with Bacillus Calmette-Guérin (BCG) [2]. However, up to 70% of these patients will experience one or more intravesical tumor recurrences, which means that cystoscopical examination is required at regular intervals to identify and remove recurrent tumors. Furthermore, 10 to 40% will eventually progress to muscle invasive bladder cancer (MIBC) and to metastatic disease. The aggressive biological behavior of MIBC coupled with limited therapeutic options results in a median survival of less than two years for patients with metastatic disease. Novel targeted treatments have the potential to inhibit the growth of recurrent NMIBC, thus reducing the burden of repeated cystectomy, and to treat MIBC, thus prolonging survival.

Fibroblast growth factors (FGF) play an important role in cellular development, wound-healing, proliferation, and angiogenesis [3]. These growth factors signal through four transmembrane glycoprotein receptors (FGFR1-4). Ligand binding leads to receptor dimerization, phosphorylation of the cytoplasmic tyrosine kinase domain and activation of downstream targets that mediate the activity of FGFs [4]. It has been recognized for some time that mutations in FGFRs, particularly FGFR3, are common in bladder cancers [5, 6]. Activating point mutations of FGFR3 are found in up to 80% of NMIBC and data from the cancer genome atlas (TCGA) suggest they can be found, along with chromosomal amplification, in 17% of MIBC as well [7]. Increased expression of wild type FGFR3 is also found in up to 40% of MIBC [8, 9]. Chromosomal translocations, including one on chromosome 4 involving FGFR3 and TACC3, have also been identified in patients [7]. These data suggest that FGFR3 is an important therapeutic target in both NMIBC and MIBC. Indeed, several studies have shown that FGFR3 inhibition has a profound inhibitory effect on some bladder cancer cell lines in preclinical models [10, 11]. Several FGFR3 inhibitors have entered clinical trials and early data is promising for several compounds [12, 13], including NVP-BGJ398 (BGJ398). In a global phase I trial, BGJ398 was found to have an acceptable adverse event profile and encouraging initial findings of efficacy in FGFR3-mutant bladder and other cancers.

BGJ398 was developed to be a highly selective FGFR inhibitor [14]. It inhibits FGFR1, FGFR2, and FGFR3 with IC_50_ ≤ 1 nM, FGFR3-K650E with IC_50_ = 4.9 nM, and FGFR4 with IC_50_ = 60 nM. Of over 70 other kinases tested, only VEGFR2 (0.18 uM), KIT (0.75 uM), and LYN (0.3 uM) were inhibited at submicromolar concentrations, demonstrating its high selectivity. Like the small molecule FGFR inhibitors PD173074, TKI-258, and SU5402, BGJ398 was shown to inhibit the growth of a subset of bladder cancer cell lines, including SW780, RT-112, and RT-4 cells [10, 14]. These cells have increased expression of non-point-mutated FGFR3 and do not show high-level gene amplification, and were much more sensitive to FGFR inhibition than cell lines with point mutations [10]. RT-112 cells have been shown to require FGFR3 activity for proliferation *in vitro* and as xenografts in mice [10, 11]. Seeking an explanation for the great sensitivity of these cell lines to FGFR3 inhibition, Williams, *et al* identified two novel fusions between FGFR3 and other proteins resulting from chromosomal translocations, in patient samples and cell lines, including a FGFR3-TACC3 fusion protein in RT-112 cells [15]. This protein is highly activated and transforms NIH-3T3 cells and at least partially explains the sensitivity of this cell line to BGJ398 and other FGFR3 inhibitors. The exquisite sensitivity of RT112 cells to FGFR3 inhibition makes them an ideal cell line in which to study resistance. As such, we developed RT112 lines resistant to BGJ398 and identified potential mechanisms of resistance, which may predict resistance mechanisms in humans.

## 2. Materials and Methods

### Cells, culture conditions and reagents

RT-112 and HEK293 cells were purchased from the ATCC and were maintained in RPMI 1640 or DMEM supplemented with 10% FBS and antibiotics, respectively. Cell line authentication has been performed by the ATCC within the last two years. NVP-BGJ398 was provided by Novartis. Other chemicals were purchased from Sigma or Cayman Chemicals. In some experiments, cells were transfected with control or FGFR expression vectors (Harvard Plasmid Repository HsCD00327305 (FGFR1), HsCD00459716 (FGFR2), HsCD00462255 (FGFR3)) using Lipofectamine LTX & Plus (Thermofisher).

### RT and qPCR

Total RNA was isolated from cells using the GeneJet RNA purification kit (Thermo Scientific). The isolated RNA was then reverse-transcribed with MMLV-reverse transcriptase (Invitrogen). Relative target-gene expression was then assessed by quantitative-PCR (qPCR) with a SYBR green detection dye (Invitrogen) and Rox reference dye (Invitrogen) on the StepOne Real Time PCR System (Applied Biosystems). Using the ΔΔCt relative quantification method, target gene readouts were normalized to RPL19 and GADPH transcript levels. Experiments are the average of biological triplicates; p values were calculated using a two-tailed Student’s t test.

### RNA-seq

RNA sequencing was performed by the City of Hope Integrative Genomics core facility. cDNA synthesis and library preparation was performed using TruSeq RNA Library prep kit in accordance with the manufacturer supplied protocols. Libraries were sequenced on the Illumina Hiseq 2500 with single read 40 bp reads. The 40-bp long single-ended sequence reads were mapped to the human genome (hg19) using TopHat and the frequency of Refseq genes was counted with customized R scripts. The raw counts were then normalized using trimmed mean of M values (TMM) method and compared using Bioconductor package “edgeR”. The average coverage for each gene was calculated using the normalized read counts from “edgeR”. Differentially regulated genes were identified using one-way ANOVA with linear contrasts to calculate p-values, and genes were only considered if the false discovery rate (FDR) was < 0.25 and the absolute value of the fold change was > 2. There were over 40.2 million reads on average with greater than 90% aligned to the human genome. Gene ontology analyses were performed using Ingenuity Pathway Analysis (Qiagen).

### Cell proliferation assays

For growth curves, cells were plated at a density of approximately 20,000 cells/well in 48 well plates. The following day, medium with vehicle or drugs was added to the cells, in quadruplicate. Proliferation was determined by measuring the DNA content of the cells in each well. Every other day, the cells were fixed in 2% paraformaldehyde, followed by staining for 5min at RT with 0.2ng/mL 4’,6-diamidino-2-phenylindole (DAPI) in PBS. The cells were washed with PBS, then read on a fluorescence plate reader (FPR) using 365/439 excitation/emission wavelengths.

## 3. Results

### Creation of resistant cell lines

RT-112 cells have been shown to be very sensitive to FGFR inhibition, and were used in the original selection and testing of NVP-BGJ398 [14]. We gradually increased the concentration of BGJ398 over time and selected two cell lines that readily grew in 3uM BGJ398, a concentration which significantly inhibited the growth of the parental drug (Figure 1A). While the resistant cells do not grow as rapidly as parental cells, their proliferation is still quite rapid. Interestingly, they have a different morphology than parental cells (Figure 1B). While parental cells maintain a uniformly circular shape, many BGJ398 resistant cells take on a flattened, crescent shape, with clusters looking as if they are forming a glandular structure.

**Figure 1:**
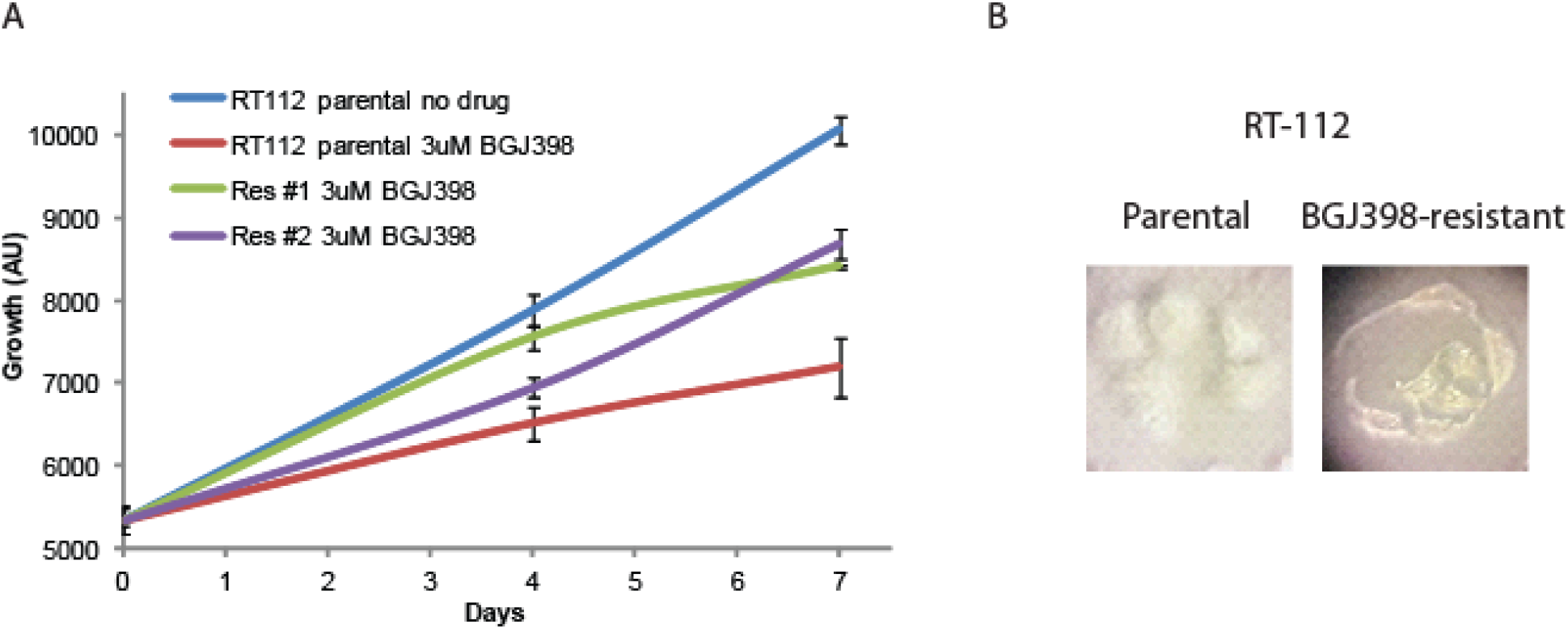
Development of NVP-BGJ398 resistant cell lines. RT-112 cells were grown in increasing amounts of NVP-BGJ398 until such time two, independent lines were able to grow in 3uM of drug. (A) Parental and resistant cell lines were plated in quadruplicate in 48-well plates and the indicated drugs were added a day 0. Cell density was measured at days 0, 4, and 7, and growth curves were created. (B) Pictures of parental and resistant lines demonstrating altered morphology.

### RNA-seq and qPCR validation

Two separate plates of parental RT-112 cells were treated with vehicle or 3uM BGJ398 overnight, at which point RNA was harvested from these cells, as well as from the two independent BGJ398 resistant cell lines that had been growing continuously in 3uM drug. To identify potential pathways of resistance, we performed RNA-sequencing and pathway analysis. Hierarchical clustering demonstrated that similar samples clustered together and that the resistant cell lines, while not having identical patterns of transcription, clustered more closely to the vehicle treated parental cells than did the drug-treated parental cells (Figure 2A). Using cut-offs described in the methods, many significant differences were found in gene regulation among the three groups (Figure 2B). The smallest number of differences was found between drug-treated parental cells and drug-resistant cells, but these are the most informative for they likely reflect the adaptive response to drug treatment. Two of the ten transcripts most decreased in the drug resistant cells were s100A8 (34 fold) and s100A9 (15 fold). However, these genes were also decreased by BGJ398 treatment in parental cells, just significantly more so in the resistant cells; this was confirmed by qPCR (Figure 2C). Interestingly, differences in several FGF-related transcripts were also significant. FGFR1 was increased in resistant cells 5 fold, while FGFR3 was decreased 4 fold. FGFR2 was increased slightly, but not significantly in the RNA-seq data. The FGF binding protein 1 (FGFBP1), which facilitates release of FGFs from the extracellular matrix [16], was decreased 2 fold. qPCR confirmed the regulation of the FGF-related factors (Figure 2D), and for each of the receptors, showed that significant changes occurred only in the resistant cell line, not in parental cells challenged with drug overnight, suggestion a unique adaptation to growth in BGJ398.

**Figure 2:**
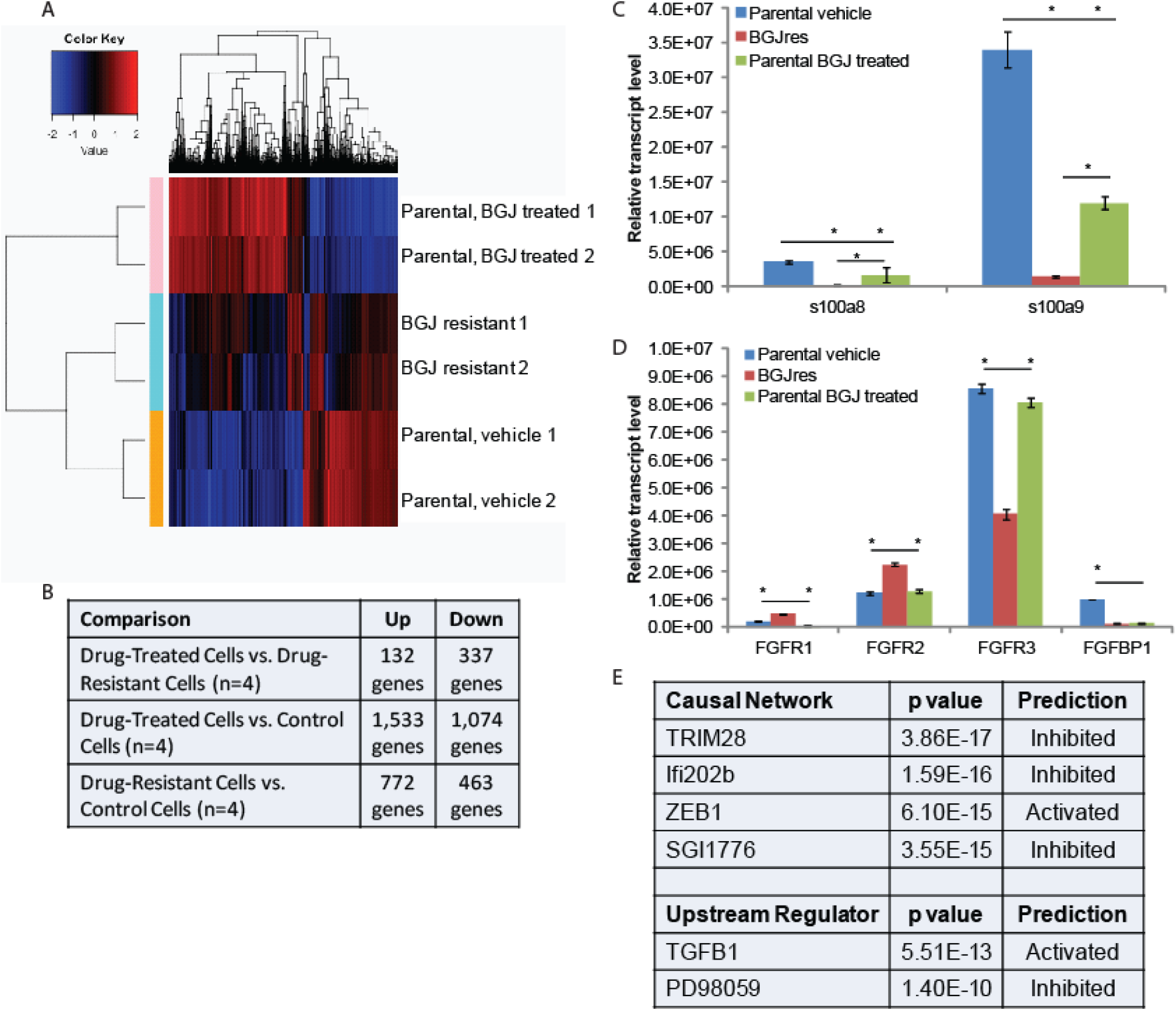
RNA-seq analysis: (A) Hierarchical clustering was performed on RNA-sequencing data from two independent untreated and BGJ398 treated parental RT-112 cell samples as well as two BGJ398 resistant RT-112 cell lines. (B) The numbers of significantly differentially regulated transcripts between treatment conditions is shown. (C,D) To validate RNA-sequencing results, RNA was extracted from the indicated cells and RT-qPCR was performed using primers for the indicated genes. S100A8 and S100A9 represent two of the most highly enriched transcripts in the drug resistant versus drug-treated parental cell datasets while the FGFR transcripts, which could play a direct role in mediating BGJ398 resistance, were also confirmed to be significantly different among treatment groups. (E) Ingenuity Pathway Analysis was performed to identify pathway signatures that were significantly different between drug resistant and drug-treated parental data sets. The most significantly different Causal Network and Upstream Regulator signatures are shown. (* p<0.05)

Ingenuity Pathway Analysis was used to identify pathways that were uniquely affected in the BGJ398 resistant cells (Figure 2E). Causal Network analysis suggested that TRIM28 (or KAP1) and Ifi202b networks, both of which regulate the interferon response [17, 18], were inhibited in BGJ398-resistant cells. Related to this, Upstream Regulatory analysis suggested that TGFβ signaling, which can repress the interferon response [19], was activated in resistant cells. Upstream regulator analysis also suggested that the pathway controlled by PD98059, a MEK1 inhibitor [20], was inhibited, which perhaps suggests that the MEK pathway is activated in resistant cells. Likewise, Causal Network analysis suggests that the pathway controlled by SGI1776, a PIM kinase inhibitor [21], was inhibited, which perhaps suggests that PIM kinases are activated in resistant cells. Finally, IPA Causal Network analysis also found the ZEB1 network to be activated in resistant cells. ZEB1 represses E-cadherin expression, driving epithelial mesenchymal transition (EMT) [22].

### FGFR3 and PIM kinase mediate resistance

To determine if changes in FGFR levels affected the growth of RT-112 cells or their sensitivity to BGJ398, we transfected FGFR1 or FGFR2 expression plasmids into parental RT-112 cells or FGFR3 into BGJ398 resistant cells and performed growth assays (Figure 3). Transient expression of FGFR1 or FGFR2 alone did not affect the growth of parental RT-112 cells, nor did it affect their sensitivity to BGJ398. However, expression of FGFR3 in BGJ398 resistant cells caused decreased growth in the presence of BGJ398. This might imply a restoration of sensitivity to the drug in these cells.

**Figure 3:**
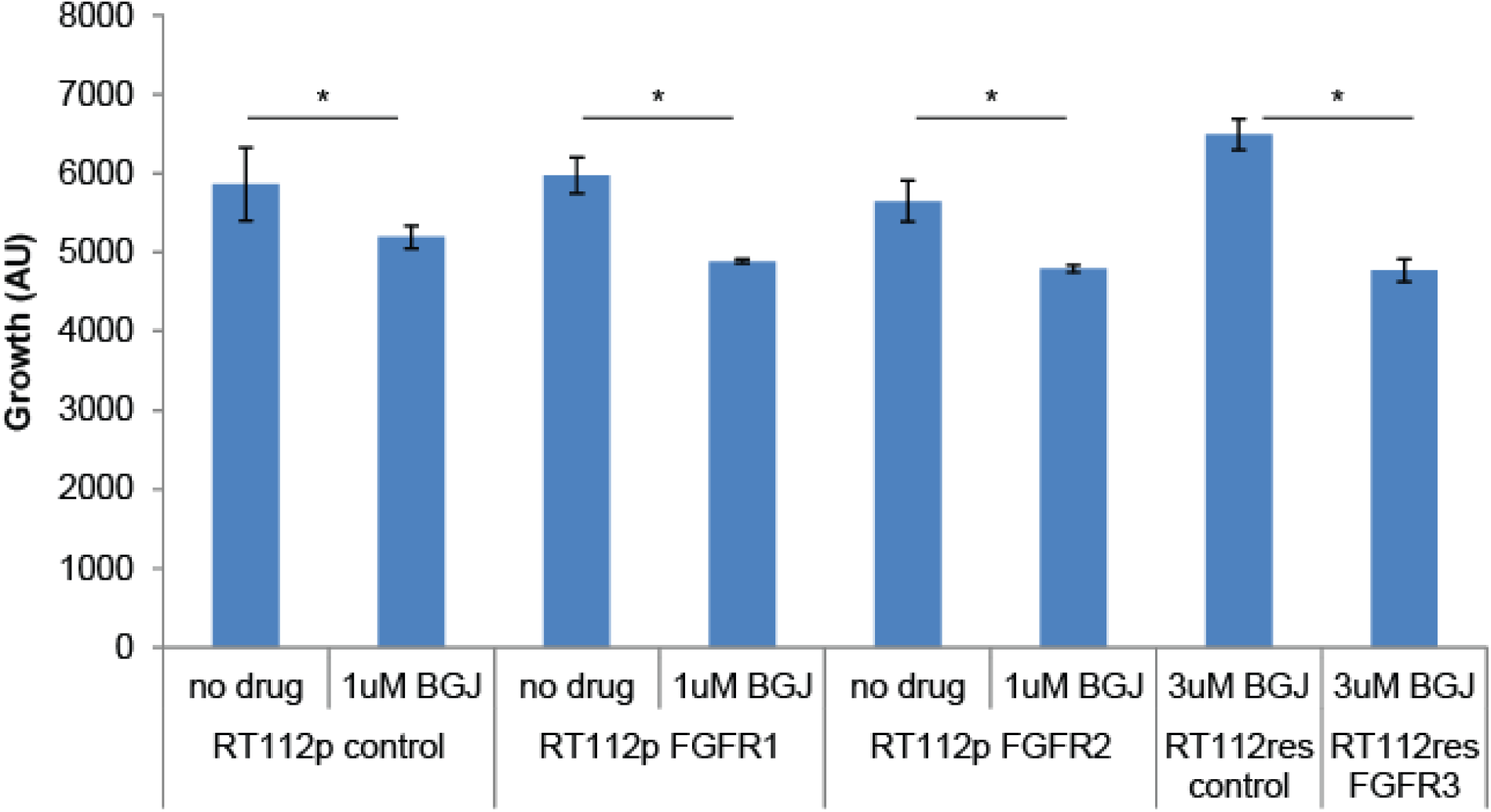
FGFR over-expression in RT-112 cells: Parental RT-112 cells were transiently transfected with a control plasmid or FGFR1 or FGFR2 expression plasmids while BGJ398 resistant cells were transfected with control or FGFR3 expression plasmids. Growth was measured over seven days by DAPI staining, and the relative cell number at day 7 is shown (AU = arbitrary units, * p<0.05).

Because the SGI1776 signal was found to be decreased in BGJ398 resistant cells, we examined what effect this inhibitor would have on the growth of parental and resistant RT-112 cells. We did not observe any significant differences in PIM transcript expression in drug treated or drug resistant cell lines (Figure 4A). However, treatment of BGJ398 resistant cells with SGI1776 significantly inhibited the growth of these cells (Figure 4B). Interestingly, a combination of BGJ398 and SGI1776 was significantly better at preventing growth of parental RT-112 cells than BGJ398 alone. SGI1776 displayed no overt toxicity at the concentration used to inhibit RT-112 cell growth as it did not inhibit the growth of HEK293 cells (Figure 4B).

**Figure 4:**
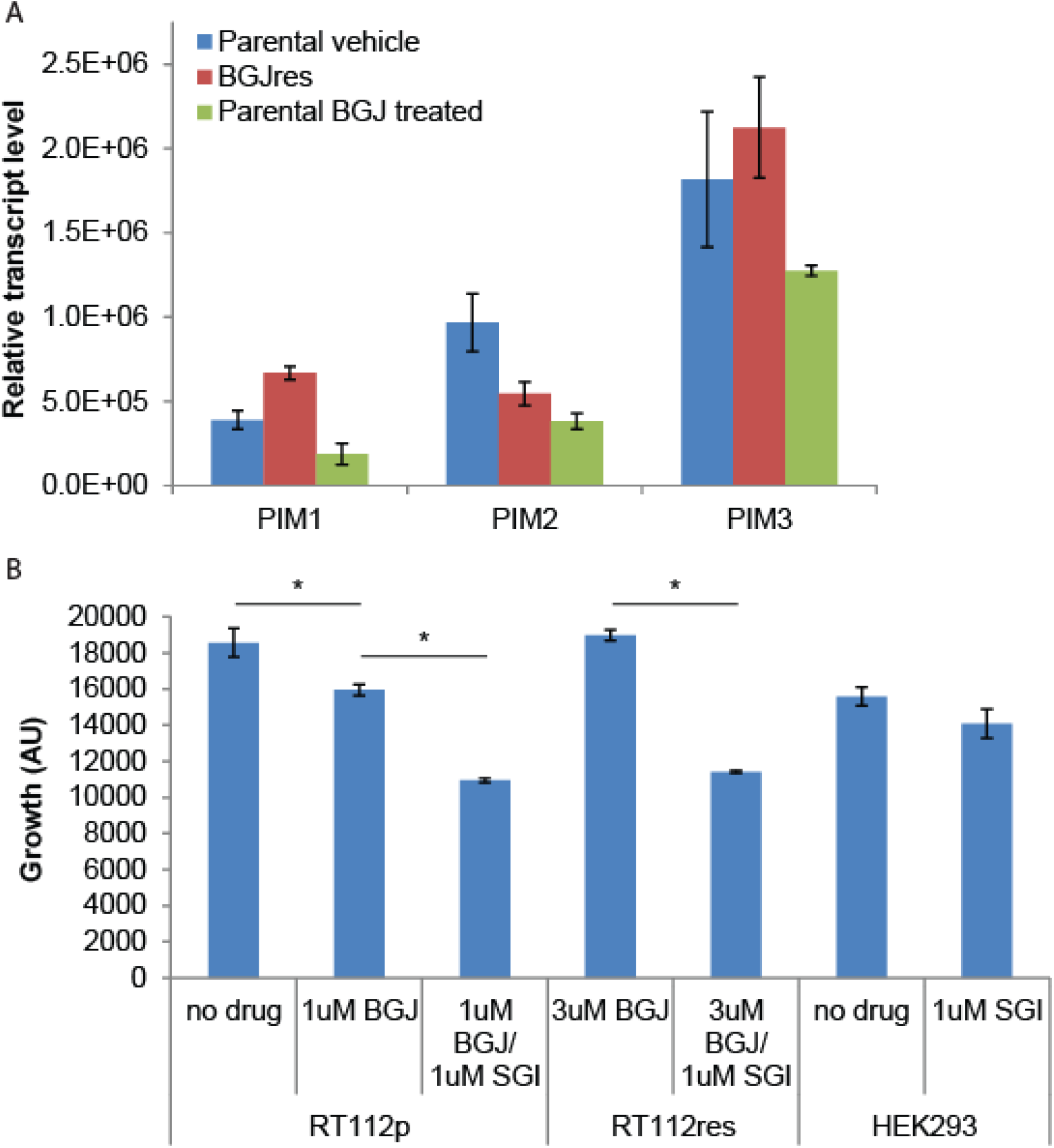
SGI1776 treatment of RT-112 cells: (A) RNA was extracted from the indicated cells and RT-qPCR was performed using primers for the three PIM transcripts. No significant differences were observed. (B) Parental and BGJ398 RT-112 cells as well as control HEK293 cells were treated with BGJ398 and/or SGI1776 as indicated and growth at day 7 is shown (AU = arbitrary units, * p<0.05).

## 4. Discussion

Recently reported data from Phase I clinical trials with two FGFR-targeted agents are very encouraging [12, 13]. Furthermore, alterations in FGFR, including FGFR3 mutations, FGFR3-TACC3 translocations, and FGFR2 alterations have been associated with response to the FGFR inhibitors JNJ-42756493 and BGJ398. Other FGFR inhibitors, including LY2874455, BMS-582664, BIBF 112, and BAY1163877 are in development for bladder and other cancers with FGFR alterations [23]. FGFR-targeting agents will hopefully soon be approved for use in bladder and other cancers, but like most targeted agents, resistance is expected to develop. Using the RT-112 cell model, which was used in the original development of BGJ398, we developed two independent resistant cell lines, which grew in 3uM of drug, well above its IC_50_ in parental cells. Using an RNA-seq approach, we identified several pathways that potentially mediate resistance to BGJ-398. Two of the most promising are alternate FGFR usage and activation of PIM kinase signaling.

We found that FGFR1 and FGFR2 transcript levels were increased in resistant cells while FGFR3 levels were decreased. BGJ398 has nearly equal affinity for all three receptors so differential receptor affinity cannot explain resistance to BGJ398. However, FGFR1 versus FGFR3 expression on bladder cancer cells is indicative of an altered phenotype and has been shown to mediate BGJ398 sensitivity [24]. Chen, *et al* found that FGFR1 was expressed on bladder cancer cells that also expressed the mesenchymal markers ZEB1 and vimentin, whereas FGFR3 expression was restricted to the E-cadherin- and p63-positive epithelial subset. Sensitivity to the growth-inhibitory effects of BGJ398 was also restricted to the epithelial cells and it correlated directly with FGFR3 mRNA levels but not with the presence of activating FGFR3 mutations. In contrast, BGJ398 did not strongly inhibit proliferation but did block invasion in the mesenchymal type bladder cancer cells *in vitro* [24]. We observed a morphological change in our BGJ398 resistant cells that could very well be a reflection of a more mesenchymal state. Indeed, in our RNA-seq data vimentin levels were increased 2.5 fold (p=0.003) in resistant cells, as were ZEB1 levels, although not significantly (p=0.11). Furthermore, our pathway analysis suggested that ZEB1 signaling was activated in resistant cells. ZEB1 is a documented mediator of epithelial-mesenchymal transition and is known to be induced by FGF2 signaling [25], further supporting a role for altered FGFR expression in lineage transition and drug resistance. In line with the Chen, *et al* report, we saw that BGJ398 sensitivity correlated with FGFR3 levels, as over-expression re-sensitized the cells to growth inhibition by BGJ398 (Figure 3). Our data, combined with the Chen, *et al* report, strongly suggest that altered FGFR expression drives, or is at least reflects a lineage transition that mediates resistance to BGJ398. This is reminiscent of the lineage plasticity that mediates anti-androgen resistance in metastatic prostate cancer [26], and may represent a wider mechanism of resistance to targeted agents. Whether lineage plasticity mediates resistance to BGJ398 and other FGFR inhibitors and exactly how such plasticity develops should be further investigated.

We also found that the PIM kinase inhibitor, SGI1776, significantly inhibits the growth of BGJ398 resistant cells. Pim1 is a serine-threonine kinase which promotes early transformation, cell proliferation, and cell survival during tumorigenesis in several cancer types, including bladder cancer, where it was found to be over-expressed in invasive cancers compared to non-invasive cancers and normal tissues [27]. Another study found high levels of expression of all three PIM family members in both non-invasive and invasive urothelial carcinomas compared to normal tissue [28]. Furthermore Pim1 knock-down [27] or treatment with the PIM kinase inhibitor TP-3654 [28] reduced the growth of several bladder cancer cell lines in culture and in xenografts. Our data that demonstrated inhibition of both parental and BGJ398 resistant cell lines using a PIM kinase inhibitor, suggesting that PIM kinase likely remains a viable target in bladder cancer, even after FGFR inhibitor resistance develops.

Pathway analysis suggested other possible mechanisms of resistance, each of which bears further investigation. Upstream regulator analysis suggested that TGFβ signaling was activated in resistant cells. TGFβ has long been known to play an important role in bladder cancer, in part through its regulation of interferons [29]. Interestingly, the two most significantly inhibited Causal Networks in the IPA analysis were those controlled by TRIM28 (or KAP1) and Ifi202b (Fig 2C), both of which also regulate the interferon response [17, 18]. S100A8 and S100A9, two of the most inhibited genes in the resistant cells, are known to be inhibited by TGFβ and activated by interferons (in part through TRIM28 and Ifi202b) [30, 31], which fits well with the IPA analysis and strongly suggests that TGFβ activation and interferon suppression is important for growth of RT-112 cells in BGJ398. TGFβ and interferons play a complicated role in tumor development and progression, making their value as therapeutic targets questionable. Upstream regulator also suggested that the pathway controlled by PD98059, a specific MEK1 inhibitor, was inhibited, which perhaps suggests that the MEK pathway is activated in resistant cells. Activation of MEK1 has been previously reported in bladder cancer and PD98059 has been shown to reduce proliferation in bladder cancer cells in vitro [32]. This suggests that MEK1 inhibition might be useful in FGFR inhibitor resistant bladder cancer as well.

In a recent report, a RT-112 cell line was developed that had resistance to the FGFR inhibitor AZD4547 [33].The authors performed a synthetic lethality RNAi screen to identify kinases that, when depleted, increased the activity of AZD4547. They identified multiple members of the phosphoinositide 3-kinase (PI3K) pathway and found that the PI3K inhibitor BKM120 acted synergistically with inhibition of FGFR in multiple cancer cell lines having FGFR mutations. Synergy was attributed to PI3K-protein kinase B pathway activity resulting from epidermal growth factor receptor or Erb-B2 receptor tyrosine kinase 3 reactivation caused by FGFR inhibition. These pathways were not identified by our transcriptomic analysis. This could be due to a difference in approach, or it could suggest that the PI3K signaling pathway is more important in mediating the response to AZD4547 than it is for NVP-BGJ398. Regardless, there are likely multiple pathways that can lead to FGFR inhibitor resistance, and each of these reports supports studies in humans to determine if these, or other, mechanisms mediate resistance in actual patients.

## 5. Conclusions

Our results suggest that altered FGFR expression and PIM kinase activity could mediate resistance to NVP-BGJ398. These pathways should be investigated in samples from patients resistant to this drug.

## 6. Acknowledgments

The authors would like to thank the Integrative Genomics core staff for assistance with this project.

## 7. Funding

Research reported in this publication included work performed in core facilities supported by the National Cancer Institute of the National Institutes of Health under award number P30CA033572. The content is solely the responsibility of the authors and does not necessarily represent the official views of the National Institutes of Health.

## 8. Author contributions

SKP helped with the design of the study, acquisition of materials, and writing of the paper. MH carried out experiments and edited the paper. JOJ managed the project, assisted with experimental design and execution, and writing of the paper

## 9. Competing interests

SKP is a consultant for Novartis. JOJ and MH have no conflicts to report.

